# pTAC3 and pTAC14 are required for binding of plastid-encoded RNA polymerase to DNA

**DOI:** 10.1101/2024.03.01.582956

**Authors:** Joyful Wang, V. Miguel Palomar, Ji-Hee Min, Andrzej T. Wierzbicki

## Abstract

Plastid-encoded RNA polymerase (PEP) is a bacterial-type multisubunit RNA polymerase responsible for the majority of transcription in chloroplasts. PEP consists of four core subunits, which are orthologs of their cyanobacterial counterparts. In *Arabidopsis thaliana*, PEP associates with 12 PEP-associated proteins (PAPs), which serve as peripheral subunits of the RNA polymerase. The exact contributions of PAPs to PEP function are still poorly understood. We use ptChIP-seq to show that PAP1/pTAC3, a peripheral subunit of PEP, binds to the same genomic loci as RpoB, a core subunit of PEP. The *pap1*/*ptac3* mutant shows a complete loss of RpoB binding to DNA throughout the genome, indicating that PAP1/pTAC3 is necessary for RpoB binding to DNA. A similar loss of RpoB binding to DNA is observed in the *pap7*/*ptac14* mutant, which is defective in another peripheral PEP subunit. We propose that the peripheral subunits of PEP are required for the recruitment of core PEP subunits to DNA.

**KEY MESSAGE:** The peripheral subunits of plastid-encoded RNA polymerase play a crucial role in recruiting the core PEP subunits to DNA in Arabidopsis chloroplasts.

## INTRODUCTION

The chloroplast genome is transcribed by two classes of RNA polymerases: nuclear-encoded RNA polymerase (NEP) and plastid-encoded RNA polymerase (PEP). NEP is a single subunit enzyme similar to T3/T7 phage RNA polymerases, which is encoded in the nuclear genome, translated by cytosolic ribosomes, and imported into plastids. NEP is primarily active in the early stages of chloroplast development. PEP, on the other hand, is a multi-subunit protein complex similar to bacterial RNA polymerases. The chloroplast genome encodes the core subunits of PEP, which is responsible for the majority of transcription in mature chloroplasts (Pfannschmidt et al. 2015).

The core PEP enzyme is composed of two α-subunits (RpoA), the catalytic β-subunit (RpoB), a β’-subunit (RpoC1) and a β’’-subunit (RpoC2) (Pfannschmidt et al. 2015). The PEP complex also interacts with nuclear-encoded sigma factors, which are responsible for promoter recognition and sequence-specific initiation of transcription (Chi et al. 2015). Although the PEP core complex has structural similarities to bacterial RNA polymerases, it can only be detected in its pure form in non-photosynthetic plastids. In chloroplasts, PEP is present in a much larger protein complex, that contains several additional subunits (Pfalz and Pfannschmidt 2013).

Peripheral subunits of PEP are also referred to as PEP-associated proteins (PAPs). They have been identified primarily through their physical interaction with the PEP complex (Pfannschmidt et al. 2000; Suzuki et al. 2004; Pfalz et al. 2006; Steiner et al. 2011). *Arabidopsis thaliana* has 12 PAPs, which are of eukaryotic origin and are encoded in the nuclear genome. The molecular architecture of the PEP complex (Ruedas et al. 2022) and functions of individual PAPs remain only partially understood (Pfannschmidt et al. 2015).

PAP proteins are necessary for the proper expression of chloroplast-encoded genes and the development of photosynthetic chloroplasts (Pfalz et al. 2006; Garcia et al. 2008; Arsova et al. 2010; Yagi et al. 2012; Gilkerson et al. 2012; Grübler et al. 2017; Yu et al. 2018; Liebers et al. 2020). The only exceptions are PAP4/FSD3 and PAP9/FSD2, which have weaker phenotypes due to partial redundancy (Myouga et al. 2008). It has been proposed that all PAPs are necessary for the proper functioning of the PEP complex, which explains why most PAP mutants exhibit similar non-photosynthetic phenotypes (Pfannschmidt et al. 2015).

The proposed requirement of all PAPs for PEP transcription predicts that PAPs should co-localize with core PEP subunits throughout the chloroplast genome. This has been partially demonstrated in wheat for PAP1/pTAC3 on a few individual loci (Yagi et al. 2012). It was found that both RpoA and PAP1/pTAC3 bind to the promoters of PEP-transcribed genes *psbA, rbcL, psaA, rrn, psbD* and *trnE* (Yagi et al. 2012). On *psaA* and *rrn*, binding signal levels of RpoA and PAP1/pTAC3 were very similar while on the remaining tested loci, PAP1/pTAC3 binding was substantially stronger than binding of RpoA (Yagi et al. 2012). It is currently unknown if PAP1/pTAC3 overlaps with the binding of core PEP subunits throughout the rest of the chloroplast genome.

The requirement of PAPs for proper expression of plastid genes predicts that in the absence of the complete set of PAPs, PEP should be unable to transcribe. However, posttranscriptional regulation plays a significant role in chloroplast gene regulation (Barkan 2011), indicating that defective transcription is not the sole explanation for PAP mutant phenotypes. Furthermore, even if PAPs are required for transcription, it remains unknown if they are required for recruitment of PEP to DNA or other aspects of PEP function.

To determine the impact of PAPs on transcription, we examined the DNA binding patterns of PAP1/pTAC3 and RpoB. Our analysis revealed that PAP1/pTAC3 binds to the same genomic regions as RpoB with comparable intensities. We subsequently investigated whether PAP1/pTAC3 is necessary for RpoB binding to DNA. The *pap1*/*ptac3* mutant exhibited no detectable RpoB binding to DNA, indicating that PAP1/pTAC3 is essential for the recruitment of PEP to its target genes. We also tested whether another peripheral PEP subunit, PAP7/pTAC14, is necessary for PEP binding to DNA. Similarly to the impact of PAP1/pTAC3, the *pap7*/*ptac14* mutant lost all detectable RpoB binding to DNA. This suggests that the requirement for PEP binding to DNA may be a more general property of PAPs.

## MATERIALS AND METHODS

### Plant materials and growth conditions

Wild-type *Arabidopsis thaliana* ecotype Col-0 was used in all analyses. We used the following mutant genotypes: *ptac3* (Salk_108852) (Alonso et al. 2003) and *ptac14* (SAIL_566_F06) (McElver et al. 2001). Experiments were performed on 5-day-old plants. Seeds were first stratified in darkness at 4°C for 48h and grown on 0.5X MS plates (0.215% MS salts, 0.05% MES-KOH pH 5.7, 1% sucrose, 0.65% agar) for 5 days at 22°C under constant white LED light (50 μmol m^-2^ sec^-1^).

### Chloroplast crosslinking

As previously described (Palomar et al. 2022), whole 5-day-old seeds were vacuum-infiltrated with 4% formaldehyde for 10 minutes and incubated in darkness for 4 h at 4°C. Formaldehyde was quenched by adding 2 M glycine to 125 mM final concentration and vacuum infiltrating for 5 minutes.

### ptChIP-seq

Crosslinked whole 5-day-old seedlings were homogenized in ice cold chloroplast lysis buffer (50 mM Tris-HCl [pH 8.0], 10 mM EDTA, 1% SDS) then filtered through 2 layers of Miracloth by centrifuging at 1500 x g for 10 sec to remove debris. Samples were sonicated to achieve DNA fragments ranging from 200 nt to 300 nt using a QSonica Q700 sonicator. The fragmented samples were incubated overnight with 5 μg of polyclonal anti-RpoB antibody (PhytoAB, San Jose, CA, USA; catalog number PHY1239) or anti-pTAC3 antibody (PhytoAB, San Jose, CA, USA; catalog number PHY0391A) and with 60 μL Protein A Dynabeads (Thermo Fisher Scientific, Waltham, MA, USA; catalog number 10002D). After incubation, the beads were washed, and DNA was eluted and reverse crosslinked as described (Rowley et al. 2013). High-throughput sequencing libraries were prepared as reported (Bowman et al. 2013) and sequenced using an Illumina NovaSeq 6000 S4 flow-cell with 150 × 150 paired-end configuration at the University of Michigan Advanced Genomics Core.

### Data analysis

The obtained raw sequencing reads were trimmed using trim_galore v.0.4.1 and mapped to the TAIR10 Arabidopsis plastid genome (www.arabidopsis.org) using Bowtie2 v.2.4.5 (Langmead and Salzberg 2012). Read counts on defined genomic regions were determined using samtools v.1.15.1 and bedtools v.2.30.0 (Quinlan and Hall 2010). ptChIP-seq signals on annotated genes were calculated by dividing reads per million (RPM)-normalized read counts from anti-RpoB, anti-pTAC3, or anti-pTAC14 ptChIP-seq by RPM-normalized read counts from input samples. ptChIP-seq enrichments on annotated genes were calculated by dividing signal levels on individual genes by the median signal level on genes in the *rpoB* operon, which is not transcribed by PEP and represents background signal levels. ptChIP-seq enrichments on genomic bins were calculated by dividing signal levels on individual bins by the signal level on the entire *rpoB* operon.

### Immunoblot analysis

To detect RpoB, RpoC1, pTAC3/PAP1and Actin in Col-0 wild type, *pap1*/*ptac3*, and *pap7*/*ptac14* mutants, total proteins were extracted by 2x SDS loading buffer (125 mM Tris-HCl, pH 6.8, 2% SDS, 0.05% Bromophenol blue, 20% glycerol, 200 mM β-mercaptoethanol). Anti-RpoB antibody (PhytoAB catalog number PHY 1239), anti-RpoC1 antibody (PhytoAB catalog number PHY0381A), anti-pTACT3/PAP1 antibody (PhytoAB catalog number PHY0391A), anti-Actin antibody (Agrisera catalog number AS13 2640), and anti-rabbit IgG antibody conjugated with horseradish peroxidase (Cell Signaling catalog number 7074) were used. Protein bands were visualized using chemiluminescence reagents (SuperSignal West Femto Maximum Sensitivity Substrate, Thermo Scientific) and a ChemiDoc Imaging System (Bio-Rad).

## RESULTS

### PAP1/pTAC3 and RpoB bind the same DNA sequences

To determine the genome-wide binding pattern of PAP1/pTAC3, we performed ptChIP-seq with an anti-pTAC3 antibody. We performed three biological replicates of the assay using Col-0 wild-type plants and used a *ptac3* knock-out mutant as a negative control. PAP1/pTAC3 binding to DNA had a complex and locus-specific pattern with a strong enrichment on rRNA genes in the inverted repeats and on several individual loci in both LSC and SSC regions (Fig. 1A). anti-pTAC3 ptChIP-seq signal was not detectable in the *ptac3* mutant negative control, which confirms the specificity of the antibody (Fig. 1A). These and other high throughput sequencing data obtained in this study may be accessed in the Plastid Genome Visualization Tool (Plavisto) at http://plavisto.mcdb.lsa.umich.edu.

**Figure 1.**
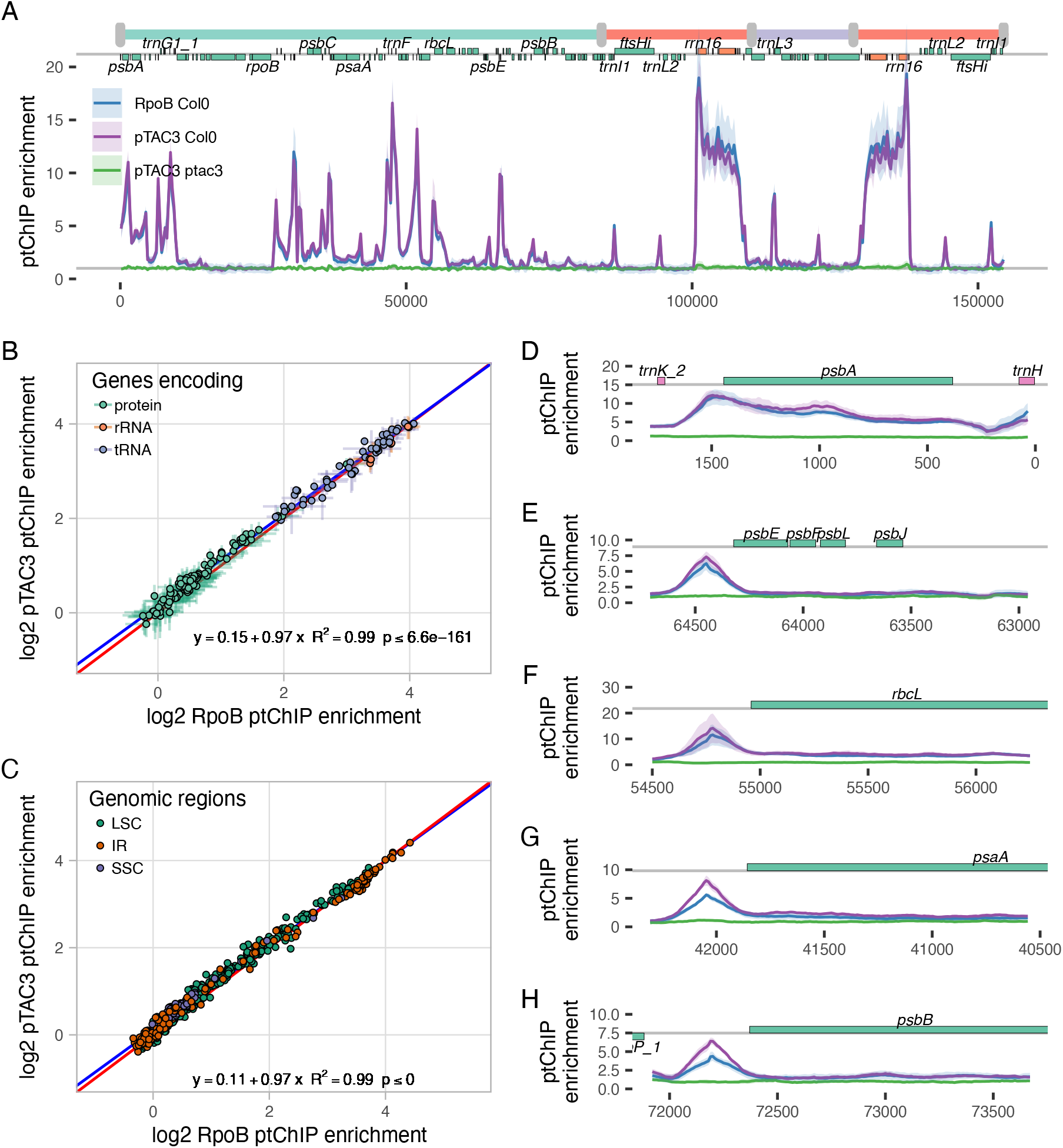
PAP1/pTAC3 and RpoB bind the same DNA sequences. A. Genome-wide patterns of pTAC3 ptChIP-seq in Col-0 wild-type and *pap1/ptac3* mutant, as well as RpoB ptChIP-seq in Col-0 wild-type. ptChIP-seq enrichment was calculated in 50-bp genomic bins and plotted throughout the entire plastid genome. Genome annotation including genomic regions, positions of annotated genes (Palomar et al. 2022), and names of selected individual genes is provided on top of the plot. Average signal from three independent biological replicates is shown. Ribbons indicate standard deviations. B. Regression analysis of PAP1/pTAC3 and RpoB binding to DNA on annotated genes. Data points are color-coded by function and show averages from three biological replicates. Error bars indicate standard deviations. The blue line represents the linear regression model. The red line represents no differences. C. Regression analysis of PAP1/pTAC3 and RpoB binding to DNA on 50-bp genomic bins. Data points are color-coded by genomic regions and show averages from three biological replicates. The blue line represents the linear regression model. The red line represents no differences. D-H. Overlap of PAP1/pTAC3 and RpoB ptChIP-seq signals on selected annotated genes. RpoB ptChIP-seq signal was calculated in 10-bp genomic bins and plotted at *psbA* (D), *psbE* (E), *rbcL* (F), *psaA* (G), and *psbB* (H) loci. Samples are color-coded as in Fig. 1A. Average signal from independent biological replicates is shown. Ribbons indicate standard deviations. Genome annotation is shown on top.

To test if PAP1/pTAC3 binds the same genomic regions as PEP core subunits, we compared the binding pattern of PAP1/pTAC3 to the binding pattern of RpoB, which was detected using three biological replicates of ptChIP-seq with anti-RpoB antibody in Col-0 wild-type seedlings. Specificity of the anti-RpoB antibody in ptChIP has been previously established (Palomar et al. 2022). The patterns of PAP1/pTAC3 and RpoB binding were remarkably similar throughout the entire chloroplast genome (Fig. 1A). To quantify the correlation between PAP1/pTAC3 and RpoB binding, we performed regression analysis using ptChIP enrichment data calculated for all annotated genes, which demonstrated a very strong (R^2^ = 0.99) and significant correlation (Fig 1B). Similar results were obtained by comparing ptChIP enrichments in 250 bp genomic bins covering the entire genome (Fig. 1C). This indicates that PAP1/pTAC3 binds the same regions as RpoB.

We further tested if PAP1/pTAC3 binding follows PEP core subunits in preferential binding to specific elements of individual protein-coding genes, particularly binding to gene promoters (Palomar et al. 2022). PAP1/pTAC3 binding was enriched on all analyzed gene promoters (Fig. 1D-H). On *psbA, psbEFLJ*, and *rbcL* genes, PAP1/pTAC3 binding closely followed binding of RpoB (Fig. 1D-F). However, on *psaA* and *psbB* promoters, PAP1/pTAC3 enrichment signal was substantially stronger than RpoB signal, which indicates that locus-specific differences between PAP1/pTAC3 and RpoB binding within individual loci are possible. Overall, these results indicate that PAP1/pTAC3 binds the same genomic regions as core subunits of PEP. This is consistent with PAP1/pTAC3 working as an accessory subunit of PEP throughout the entire chloroplast genome.

### PAP1/pTAC3 is required for RpoB binding to DNA

PAP1/pTAC3 has been proposed to work as an accessory subunit of PEP and have an impact on the accumulation of *psaA, psbA* and *rbcL* mRNAs in wheat and rice (Yagi et al. 2012; Wang et al. 2016). However, the impact of PAP1/pTAC3 on PEP transcription and especially on the recruitment of core subunits of PEP to transcribed genes remains unknown. To test if PAP1/pTAC3 is required for PEP binding to DNA, we performed ptChIP-seq with the anti-RpoB antibody in Col-0 wild-type and *ptac3* knock out mutant plants. While RpoB binding was detected at the expected loci (Palomar et al. 2022) in Col-0 wild type, no substantial binding of RpoB to DNA was detected in the *ptac3* mutant (Fig. 2A). This indicates that PAP1/pTAC3 is required for RpoB binding to DNA throughout the entire chloroplast genome.

**Figure 2.**
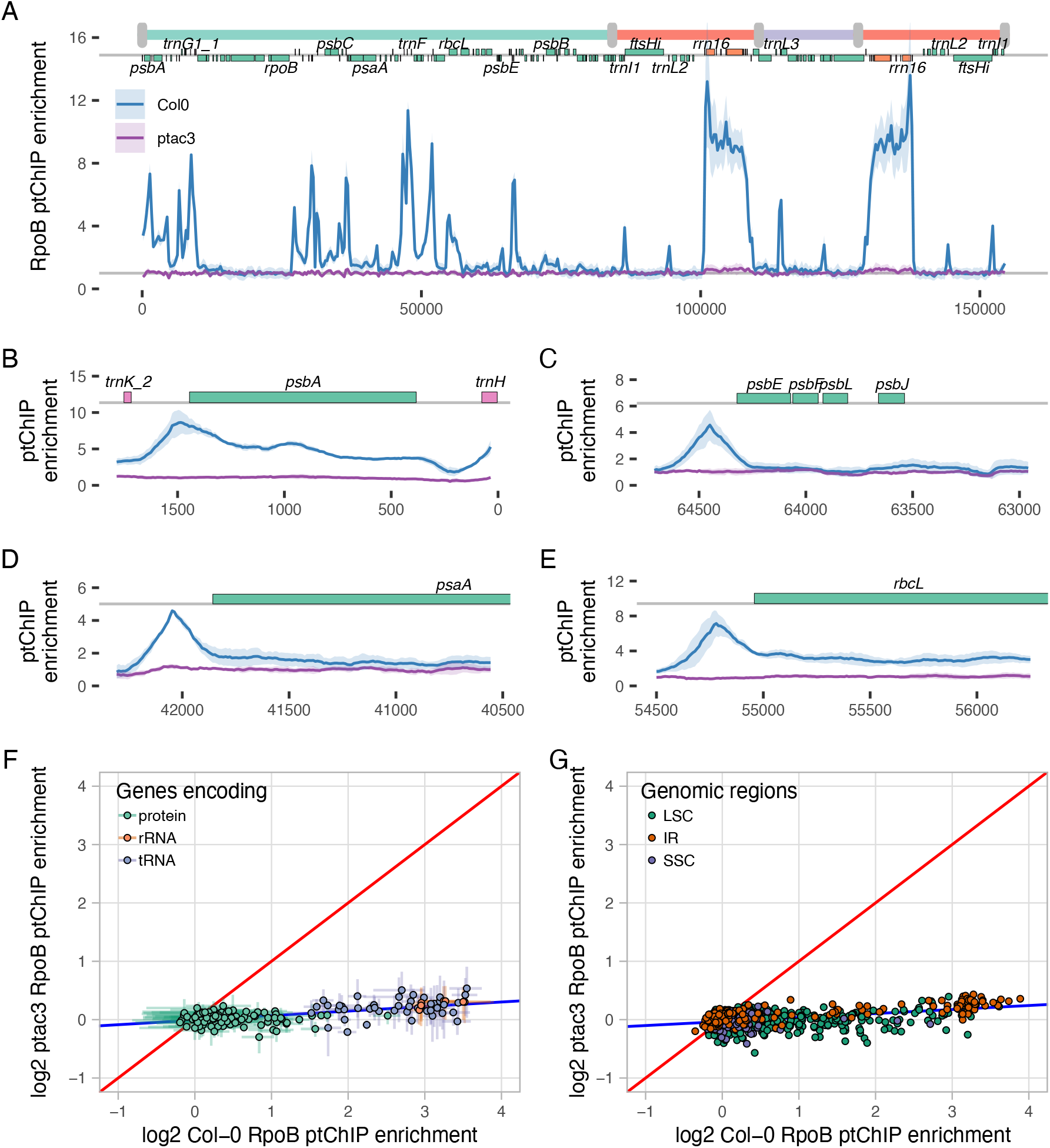
PAP1/pTAC3 is required for RpoB binding to DNA. A. Genome-wide patterns of RpoB ptChIP-seq in Col-0 wild-type and the *pap1/ptac3* mutant. ptChIP-seq enrichment was calculated in 50-bp genomic bins and plotted throughout the entire plastid genome. Genome annotation including genomic regions, positions of annotated genes (Palomar et al. 2022), and names of selected individual genes is provided on top of the plot. Average signal from three independent biological replicates is shown. Ribbons indicate standard deviations. B-E. RpoB ptChIP-seq signals in Col-0 wild-type and *pap1/ptac3* mutant on selected annotated genes. RpoB ptChIP-seq signal was calculated in 10-bp genomic bins and plotted at *psbA* (B), *psbE* (C), *psaA* (D), and *rbcL* (E) loci. Samples are color-coded as in Fig. 2A. Average signal from independent biological replicates is shown. Ribbons indicate standard deviations. Genome annotation is shown on top. F. Regression analysis of RpoB binding to DNA in Col-0 wild-type and *pap1/ptac3* mutant on annotated genes. Data points are color-coded by genomic regions and show averages from three biological replicates. Error bars indicate standard deviations. The blue line represents the linear regression model. The red line represents no differences. G. Regression analysis of RpoB binding to DNA in Col-0 wild-type and *pap1/ptac3* mutant on 50-bp genomic bins. Data points are color-coded by genomic regions and show averages from three biological replicates. The blue line represents the linear regression model. The red line represents no differences.

RpoB binding to DNA was lost in the *ptac3* mutant throughout the entire lengths of the analyzed genes, including *psbA, psbEFLJ, psaA* and *rbcL* (Fig. 2B-E). This includes the loss of RpoB binding to gene promoters where RpoB is normally enriched (Palomar et al. 2022). This is consistent with PAP1/pTAC3 being required for both initiation and elongation of PEP transcription.

Regression analysis further supports the genome-wide loss of RpoB binding to DNA in the *ptac3* mutant on annotated genes (Fig. 2F) and on 250 bp bins distributed throughout the entire genome (Fig. 2G). Small residual RpoB ptChIP enrichment signal may be observed in *ptac3* on rRNA and tRNA genes (Fig. 2F). This low signal is unlikely to be specific and may be the outcome of sequencing bias caused by differences in CG content. Overall, we conclude that PAP1/pTAC3 is required for binding of core PEP subunits to DNA throughout the entire genome.

### PAP1/pTAC3 contributes to efficient recruitment of core subunits to DNA

The loss of RpoB binding to DNA in the *pap1*/*ptac3* mutant may be caused by defective recruitment of PEP core subunits to DNA in the absence of PAP1/pTAC3. Alternatively, it may be caused by core subunit expression and/or stability being dependent on the presence of PAP1/pTAC3. To distinguish between these possibilities, we performed western blots with anti-RpoB and anti-RpoC1 antibodies in the *pap1*/*ptac3* mutant. The accumulation of RpoB and RpoC1 was reduced by approximately 50% and 40%, respectively (Fig. 3A). This indicates that reduced stability of the core PEP complex may contribute to the observed loss of RpoB binding to DNA in the *pap1*/*ptac3* mutant. However, this reduction alone cannot explain the complete genome-wide loss of RpoB binding to DNA in *pap1*/*ptac3* (Fig. 2A). Therefore, we conclude that the loss of RpoB binding to DNA in the *pap1*/*ptac3* mutant is at least partially caused by defective recruitment of RpoB to DNA in the absence of PAP1/pTAC3. This indicates that PAP1/pTAC3 contributes to the efficient recruitment of core PEP subunits to DNA.

**Figure 3.**
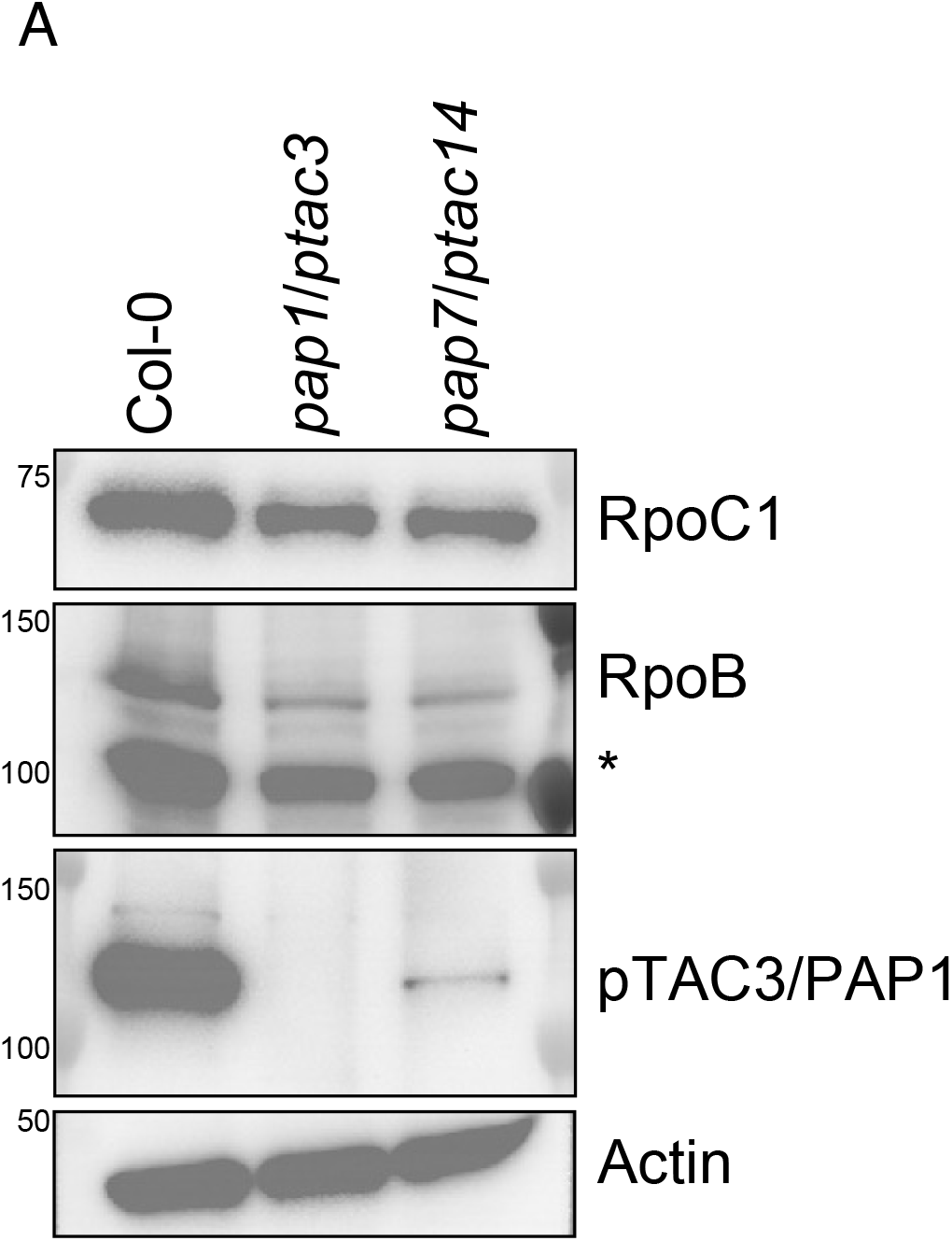
RpoB and RpoC1 are still expressed in *pap1*/*ptac3* and *pap7*/*ptac14* mutants. A. Western blot was performed using whole cell extracts from 4 day-old seedlings of Col-0 wild type, *pap1/ptac3*, and *pap7/ptac14* using anti-RpoC1, anti-RpoB and anti-pTAC3 antibodies. anti-Actin antibody was used as a loading control. Star indicates a non-specific band.

### PAP7/pTAC14 is required for RpoB binding to DNA

Requirement of PAP1/pTAC3 for PEP binding to DNA may indicate that peripheral subunits of PEP may be generally required to recruit PEP to DNA. This would be consistent with the observed reductions of PEP transcripts accumulation in most mutants defective in PAPs (Pfalz et al. 2006; Garcia et al. 2008; Arsova et al. 2010; Yagi et al. 2012; Gilkerson et al. 2012; Grübler et al. 2017; Yu et al. 2018; Liebers et al. 2020). To test this possibility, we performed ptChIP-seq with anti-RpoB antibody in a mutant defective in another accessory PEP subunit, PAP7/pTAC14. Compared to Col-0 wild type, the *pap7*/*ptac14* mutant lost RpoB binding to DNA throughout the entire chloroplast genome (Fig. 4A). Loss of RpoB binding to DNA was observed throughout the entire lengths of *psbA, psbEFLJ, psaA* and *rbcL* genes, including promoters and coding sequences (Fig. 4B-E). Very little residual signal was observed in the pap7/*ptac14* mutant on annotated genes (Fig. 4F) and on 250 bp bins distributed throughout the entire genome (Fig. 4G).

**Figure 4.**
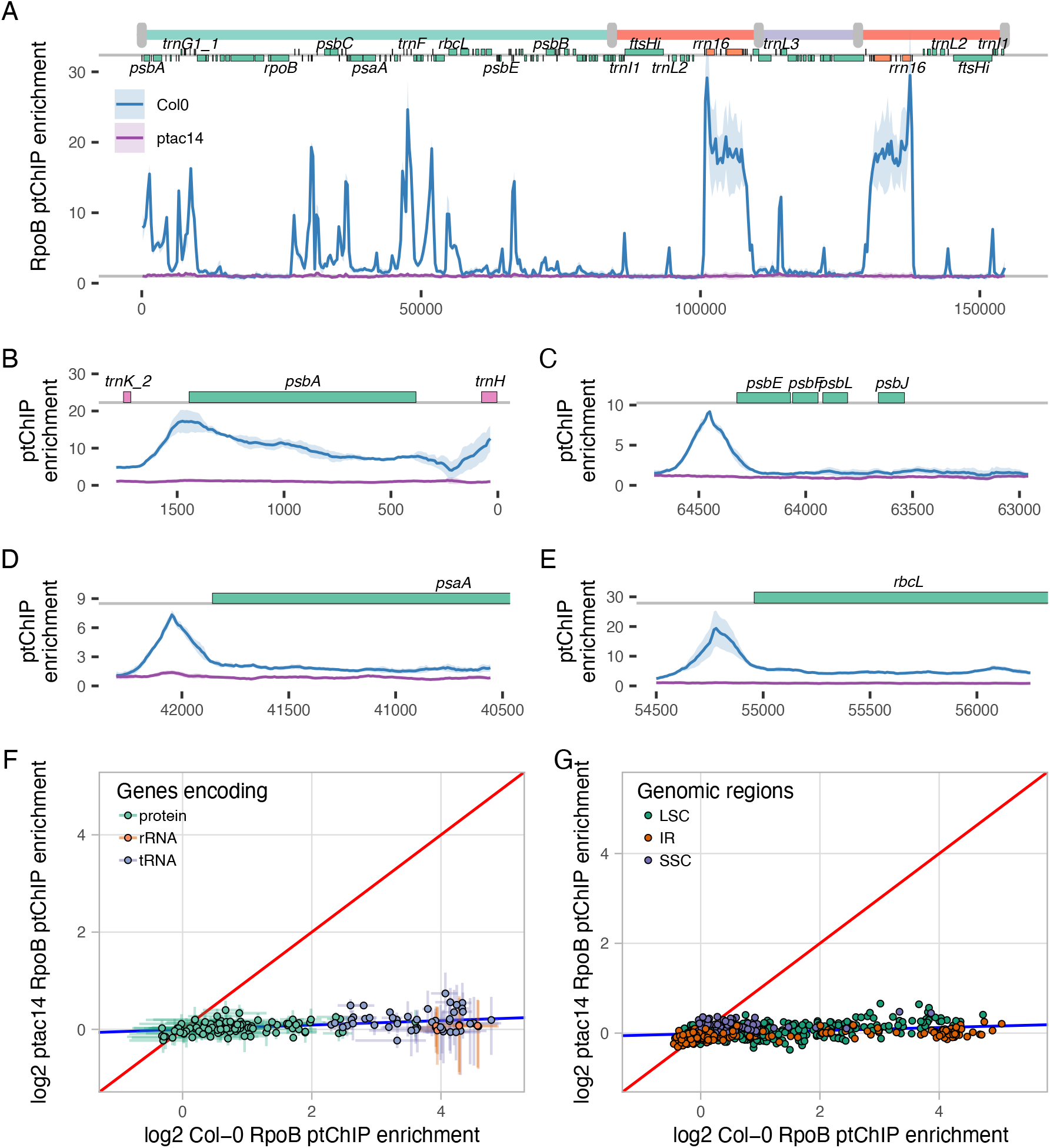
PAP7/pTAC14 is required for RpoB binding to DNA. A. Genome-wide patterns of RpoB ptChIP-seq in Col-0 wild-type and the *pap7/ptac14* mutant. ptChIP-seq enrichment was calculated in 50-bp genomic bins and plotted throughout the entire plastid genome. Genome annotation including genomic regions, positions of annotated genes (Palomar et al. 2022), and names of selected individual genes is provided on top of the plot. Average signal from three independent biological replicates is shown. Ribbons indicate standard deviations. B-E. RpoB ptChIP-seq signals in Col-0 wild-type and *pap7/ptac14* mutant on selected annotated genes. RpoB ptChIP-seq signal was calculated in 10-bp genomic bins and plotted at *psbA* (B), *psbE* (C), *psaA* (D), and *rbcL* (E) loci. Samples are color-coded as in Fig. 2A. Average signal from independent biological replicates is shown. Ribbons indicate standard deviations. Genome annotation is shown on top. F. Regression analysis of RpoB binding to DNA in Col-0 wild-type and *pap7/ptac14* mutant on annotated genes. Data points are color-coded by genomic regions and show averages from three biological replicates. Error bars indicate standard deviations. The blue line represents the linear regression model. The red line represents no differences. G. Regression analysis of RpoB binding to DNA in Col-0 wild-type and *pap7/ptac17* mutant on 50-bp genomic bins. Data points are color-coded by genomic regions and show averages from three biological replicates. The blue line represents the linear regression model. The red line represents no differences.

RpoB and RpoC1 were still present in the *pap7/ptac14* mutant at levels very similar to that observed in *pap1/ptac3* (Fig. 3A). This indicates that PAP7/pTAC14 also contributes to efficient recruitment of core PEP subunits to DNA. Interestingly, accumulation of PAP1/pTAC3 was almost entirely lost in the *pap7*/*ptac14* mutant (Fig. 3A), indicating that the impact of PAP7/pTAC14 on the recruitment of core PEP subunits may be direct and/or indirect by stabilizing PAP1/pTAC3.

Overall, we conclude that the effect of PAP7/pTAC14 on PEP recruitment to DNA is very similar to that of PAP1/pTAC3. This is consistent with accessory subunits of PEP being required for the recruitment of core PEP subunits to DNA.

## DISCUSSION

Our findings offer further support for the model where all PAPs are required for proper function of the entire PEP complex (Pfannschmidt et al. 2015). The model’s first crucial element is genome-wide co-localization of core and peripheral subunits of PEP, which we demonstrated for PAP1/pTAC3 and RpoB (Fig 1). Consistently, previous studies demonstrated genome-wide co-localization of PAP5/pTAC12 with RpoB (Palomar et al. 2022) and locus-specific co-localization of PAP1/pTAC3 with RpoA (Yagi et al. 2012). It is expected that other PAPs will also co-localize with core PEP subunits, although their genomic locations have not yet been tested.

The model predicts that all PAPs are required for PEP transcription (Pfannschmidt et al. 2015). Previous studies have shown that most PAPs are necessary for proper accumulation of PEP-transcribed RNAs (Pfalz et al. 2006; Garcia et al. 2008; Arsova et al. 2010; Yagi et al. 2012; Gilkerson et al. 2012; Grübler et al. 2017; Yu et al. 2018; Liebers et al. 2020). However, it remained unknown if PAPs affect transcriptional or posttranscriptional steps of gene expression. We found that PAP1/pTAC3 and PAP7/pTAC14 contribute to PEP recruitment to DNA (Fig. 2 and 4). This indicates that both tested PAPs are essential for PEP transcription. While the effect of other PAPs on PEP transcription has not been experimentally tested, they are also expected to be required for PEP transcription.

The domain structure of PAP1/pTAC3 is consistent with DNA and/or RNA binding (Yagi et al. 2012; Pfannschmidt et al. 2015). However, both previously published data (Yagi et al. 2012) and our own ChIP data (Fig. 1) do not provide conclusive evidence that PAP1/pTAC3 directly binds to DNA. Because ChIP includes formaldehyde crosslinking, it may detect both direct and indirect protein-DNA interactions (Hoffman et al. 2015). Therefore, it remains unclear whether PAP1/pTAC3 binds to DNA directly or indirectly through other PAPs or core PEP subunits. It is also not known how PAP1/pTAC3 contributes to PEP transcription. The observed defects in the *pap1*/*ptac3* mutant likely result from the complete loss of PEP activity and do not provide information about the specific role of PAP1/pTAC3 within the PEP complex (Pfannschmidt et al. 2015).

PAP7/pTAC14 contains a SET-domain and has been proposed to function as a protein methyltransferase (Gao et al. 2011; Pfannschmidt et al. 2015). Although its enzymatic activity and potential substrates have not been identified, we demonstrate that PAP7/pTAC14 is necessary for PEP binding to DNA (Fig. 4). This likely reflects the crucial role of PAP7/pTAC14 in PEP function, which is further supported by the albino phenotype and disrupted accumulation of PEP transcripts in the *pap7*/*ptac14* mutant (Gao et al. 2011).

The mechanisms behind the requirement of PAP for PEP transcription remain unclear. It has been demonstrated that the core PEP complex, isolated from *Sinapis alba* etioplasts and lacking PAPs (peak B), is transcriptionally active *in vitro* (Pfannschmidt and Link 1994). Our data show that PAPs co-localize with core PEP subunits throughout the entire genome, including both gene promoters and coding sequences, and that both tested PAPs are necessary for PEP binding to these regions. This suggests that in mature leaf chloroplasts, PAPs are required for both initiation and elongation of transcription. It is possible that the PEP holoenzyme requires all core and peripheral subunits to properly assemble. Determination of the exact roles of PAPs in PEP transcription remains an important goal for future studies.

## ACKNOWLEDGEMENTS

This work was supported by a grant from the National Science Foundation (MCB 1934703) to A.T.W and partially by UNAM-PAPIIT grant IA203424 to V.M.P. The sequencing data from this study have been submitted to the NCBI Gene Expression Omnibus (GEO; http://www.ncbi.nlm.nih.gov/geo/) under accession number GSE259283. Sequencing data presented in this study are available through a dedicated publicly available Plastid Genome Visualization Tool (Plavisto) at http://plavisto.mcdb.lsa.umich.edu.

## FIGURE LEGENDS

**Table 1.** High throughput sequencing datasets generated in this study.

| Experiment | Genotype and Treatment | Repl. | Numbers of paired-end sequenced reads |  |  | Plastid Genome Coverage | Median Fragment Size | GEO accession |
| --- | --- | --- | --- | --- | --- | --- | --- | --- |
|  |  |  | Raw | Mapped to whole genome | Mapped to plastid genome |  |  |  |
| RpoB ChIP | Col0 (control for ptac3) ChIP | 1 | 2404550 | 1888259 | 1378496 | 2677.1 | 172 | GSM8113044 |
|  |  | 2 | 1138407 | 909536 | 461756 | 896.7 | 196 | GSM8113045 |
|  |  | 3 | 750595 | 646869 | 235536 | 457.4 | 156 | GSM8113046 |
|  | Col0 (control for ptac3) input | 1 | 936977 | 864265 | 498015 | 967.2 | 226 | GSM8113047 |
|  |  | 2 | 921259 | 852732 | 413192 | 802.4 | 197 | GSM8113048 |
|  |  | 3 | 1234488 | 1166734 | 609706 | 1184.1 | 164 | GSM8113049 |
|  | ptac3 ChIP | 1 | 976752 | 321579 | 80814 | 156.9 | 200 | GSM8113050 |
|  |  | 2 | 611286 | 442617 | 116572 | 226.4 | 196 | GSM8113051 |
|  |  | 3 | 908445 | 625126 | 175567 | 341.0 | 180 | GSM8113052 |
|  | ptac3 input | 1 | 772403 | 695333 | 344712 | 669.4 | 217 | GSM8113053 |
|  |  | 2 | 1576591 | 1150485 | 604588 | 1174.1 | 197 | GSM8113054 |
|  |  | 3 | 1100997 | 1026495 | 479496 | 931.2 | 155 | GSM8113055 |
|  | Col0 (control for ptac14) ChIP | 1 | 873695 | 220455 | 107336 | 208.4 | 163 | GSM8113056 |
|  |  | 2 | 1236307 | 725191 | 539096 | 1046.9 | 150 | GSM8113057 |
|  |  | 3 | 1897241 | 1376761 | 1102743 | 2141.6 | 165 | GSM8113058 |
|  | Col0 (control for ptac14) input | 1 | 1499182 | 927972 | 439784 | 854.1 | 190 | GSM8113059 |
|  |  | 2 | 802892 | 636973 | 357116 | 693.5 | 191 | GSM8113060 |
|  |  | 3 | 1217644 | 1063170 | 528814 | 1027.0 | 186 | GSM8113061 |
|  | ptac14 ChIP | 1 | 789424 | 255597 | 55548 | 107.9 | 192 | GSM8113062 |
|  |  | 2 | 1931132 | 677442 | 143159 | 278.0 | 161 | GSM8113063 |
|  |  | 3 | 764132 | 479414 | 130058 | 252.6 | 169 | GSM8113064 |
|  | ptac14 input | 1 | 1616121 | 1434207 | 583153 | 1132.5 | 191 | GSM8113065 |
|  |  | 2 | 1980316 | 1793825 | 739957 | 1437.0 | 172 | GSM8113066 |
|  |  | 3 | 1424172 | 1310173 | 654756 | 1271.6 | 179 | GSM8113067 |
| pTAC3 ChIP | Col0 ChIP | 1 | 907232 | 747871 | 403277 | 783.2 | 161 | GSM8113068 |
|  |  | 2 | 1593581 | 1408513 | 568131 | 1103.3 | 156 | GSM8113069 |
|  |  | 3 | 765690 | 558636 | 225969 | 438.8 | 167 | GSM8113070 |
|  | Col0 input | 1 | 727417 | 658851 | 256060 | 497.3 | 192 | GSM8113071 |
|  |  | 2 | 1030683 | 987586 | 525899 | 1021.3 | 171 | GSM8113072 |
|  |  | 3 | 871242 | 810989 | 296145 | 575.1 | 196 | GSM8113073 |
|  | ptac3 ChIP | 1 | 1097835 | 779378 | 254734 | 494.7 | 195 | GSM8113074 |
|  |  | 2 | 1260348 | 235369 | 52143 | 101.3 | 175 | GSM8113075 |
|  |  | 3 | 1596702 | 1043009 | 157847 | 306.5 | 170 | GSM8113076 |
|  | ptac3 input | 1 | 1097835 | 1038704 | 485938 | 943.7 | 171 | GSM8113077 |
|  |  | 2 | 1260348 | 1159459 | 365804 | 710.4 | 187 | GSM8113078 |
|  |  | 3 | 790264 | 734169 | 367346 | 713.4 | 179 | GSM8113079 |

